# Wild red wolf *Canis rufus* poaching risk

**DOI:** 10.1101/2020.12.08.416032

**Authors:** Suzanne W. Agan, Adrian Treves, Lisabeth Willey

**Affiliations:** Environmental Studies Department, Antioch University New England, Keene, New Hampshire, United States of America; Carnivore Coexistence Lab, Nelson Institute for Environmental Studies, University of Wisconsin, Madison, Wisconsin, United States of America.

## Abstract

The reintroduced red wolf population in northeastern North Carolina declined to 7 known wolves by October 2020. Poaching (illegal killing) is the major component of verified anthropogenic mortality in this and many other carnivore populations, but it is still not well understood. Poaching is often underestimated, partly as a result of cryptic poaching, when poachers conceal evidence. Cryptic poaching inhibits our understanding of the causes and consequences of anthropogenic mortality which is important to conservation as it can inform us about future population patterns within changing political and human landscapes. We estimate risk for marked adult red wolves of 5 causes of death (COD: legal, nonhuman, unknown, vehicle and poached) and disappearance, describe variation in COD in relation to hunting season, and compare time to disappearance or death. We include unknown fates in our risk estimates. We found that anthropogenic COD accounted for 0.724 – 0.787, including cryptic and reported poaching estimated at 0.510 – 0.635 of 508 marked animals. Risk of poaching and disappearance was significantly higher during hunting season. Mean time from collaring until nonhuman COD averaged 376 days longer than time until reported poached and 642 days longer than time until disappearance. Our estimates of risk differed from prior published estimates, as expected by accounting for unknown fates explicitly. We quantify the effects on risk for three scenarios for disappearances, which span conservative to most likely COD. Implementing proven practices that prevent poaching or hasten successful reintroduction may reverse the decline to extinction in the wild of this critically endangered population. Our findings add to a growing literature on endangered species protections and enhancing the science used to measure poaching worldwide.

## Introduction

Many large carnivores play a significant role in the function of ecosystems as keystone species through top-down regulation, resulting in increased biodiversity across trophic levels [1–4]. They do this by influencing prey populations through predation or behavioral changes of surviving prey, potentially influencing plants, scavengers, and an array of interacting species [5]. They also influence smaller predator populations, which again has effects on smaller prey species. In California, the absence of coyotes, a top predator in that ecosystem, behaviorally released opossums, foxes, and house cats which preyed heavily on song birds and decreased the number of species of scrub-dependent birds [6]. In Yellowstone, the return of wolves as a keystone species triggered top down effects that were wide-spread throughout the park [4]. If the presence of a top predator such as a wolf increases biodiversity and has positive effects on ecosystem function, then the absence of such carnivores can simplify biodiversity long-term and significantly impair ecosystem processes [7].

Red wolves are the dominant top carnivore in the only ecosystem they occur wild, are believed to have evolved only in North America, and are critically endangered [8]. Restoring them to their native ecosystems in ecologically functional numbers would meet both the explicit and implicit demands of the Endangered Species Act (ESA) [9]. The USFWS is the implementing agency for terrestrial endangered species and it followed the ESA listing of the species by reintroducing them to northeastern NC (NENC). Management includes a permanent injunction in accordance with the 10(j) rule published in 1995 prohibiting the take of red wolves except for threat to human, livestock, or pet safety [10]. However, the reintroduced red wolf population in NENC has been declining over the last 12 years to a low of only 7 known wolves in 2020 [11] and anthropogenic mortality remains the leading cause of death.

Even with legal protections, anthropogenic mortality is causing population decline in NENC red wolves and many mammalian carnivores around the world [12]. In studying losses of Mexican wolves, *Canis lupus baileyi,*, researchers found anthropogenic mortality accounted for 81% of all deaths [13]. Many studies show wolves face high levels of anthropogenic mortality [14–17], however the persistence of some carnivore populations in areas of high human density show human-carnivore coexistence with minimal killing is possible under certain conditions [18,19]. For example, Carter et al. found that tigers and humans co-existed even at small spatial scales, likely because tigers adjusted their activity to avoid encounters with people [19]. In North America, Linnel et al., found that carnivore populations increased after favorable policy was introduced (e.g., ESA protections), despite increases in human population density [20]. Finally Dickman et al, (2011) assessed how people changed their behavior, adopting non-lethal methods for coexistence through financial incentives [21]. With very small populations and limited genetic diversity, the loss of even one red wolf could affect recovery negatively [13]. For the ESA requirement to abate human-caused mortality in endangered species, those causes should be measured accurately, understood rigorously, and intervened against effectively.

The major cause of death in red wolves is poaching [22–24]. Poaching (illegal killing) is a major component of anthropogenic mortality in many carnivore populations [16,25], but it is still not well understood [26], disrupts management efforts [27], and is often under-estimated [28]. This underestimation is partly the result of cryptic poaching, when poachers conceal evidence [17,26,27,29], and it inhibits our understanding of animal life histories, policy interventions, and management actions [26,29]. As a percent of all mortality, poaching accounted for 24 – 75% for different carnivore species and areas [30,31]. More than half of wolf mortalities in Scandinavia over the last decade are the result of poaching, with 66% of those having evidence concealed by the poacher [28]. In Wisconsin, poaching accounted for 39% - 45% of all wolf mortalities over a 32 year period with an estimated 50% cryptic, but this number is believed to be an underestimate because of non-reporting and uncertainty [16,17,26,28][17]. Although the USFWS invested large amounts in intervening to stop hybridization between red wolves and coyotes, the potentially stronger negative effects of anthropogenic mortality may deserve more attention.

Anthropogenic mortality decreases mean life expectancy for wolves, which has population wide effects. Red wolves rarely live over 10 years in the wild but up to 14 years in captivity according to the USFWS [32] however, red wolves in NENC live an average of only 3.2 years in the wild with breeding pair duration of only 2 years [33]. This prevents the development of a multigenerational social structure and pack stability, a key factor in preventing hybridization with coyotes [33–35]. Theoretically, wild populations could compensate for anthropogenic mortality through decreases in natural mortality or increases in productivity [36]. However, the strength of compensation in wolf populations is an area of active scientific controversy [14,37–39]. Also, poaching can be additive or super-additive if poachers kill a breeder or a lactating female. Super-additive mortality would result if death of a breeder also led to death of offspring. When dominant male red wolves are killed, there can be an increase in offspring mortality through mate turnover [40]. This is only more prominent in small populations at low density such as red wolves in NENC. If there is a large portion of non-breeding adults in the remaining population there is potential for recovery since new breeding pairs could take up residence, however long term compensation for anthropogenic mortality depends on survival of those adults [36]. The Red Wolf Species Survival Plan and USFWS 5-year review both call for removal of threats that have the potential to bring about the extinction of red wolves [41,42]. Currently, anthropogenic mortality is the leading cause of death for wild red wolves in the endangered NENC population [22]. In the first 25 years of reintroduction, 72% of known mortalities were caused by humans, and in many cases avoidable. These included suspected illegal killing, vehicle strikes, and private trapping [24]. Gunshot mortalities alone increased by 375% in the years 2004 – 2012 compared to the 5 previous years [24] and increased 7.2 times during deer hunting season when compared to the rest of the year. A higher percentage of wolves that go missing are unrecovered during this same time as opposed to outside the hunting season [23]. Human-caused mortalities also accounted for 40.6% of all breeding pair disbandment, mostly from gunshots [23] and natural replacement has decreased since the mid-2000’s [23]. Poaching has not been adequately prevented [22]. Therefore, precise measurement of mortality using the most current and comprehensive data is critical to understand how sources of mortality influence red wolf population dynamics and its legal recovery under the ESA [22,25].

Measuring mortality risk, the proportion of all deaths attributable to a given cause, depends on many factors including the ability to monitor individuals over time. Following individuals with GPS or VHF technology is the standard method but can be costly, compromise animal welfare, and can create systematic bias when the technology fails, or wolves move out of range [43]. Because killing a red wolf is illegal under the ESA except in case of imminent harm to a human, there may be incentives for poachers to destroy evidence, including radio-collars, which limits the data available to researchers and adds to bias. Destruction of evidence is rarely, if ever, associated with nonhuman COD or legal human-induced COD [16]. Therefore, measurement error caused by cryptic poaching adds a systematic bias. The possibility that marked animals were poached and monitoring interrupted by destruction of transmitters should be considered in poaching estimates lest we under-estimate the risk of poaching by considering only the subset that is observed by officials because the poacher left a transmitter intact. Problems such as these can be addressed with models that allow multiple sources of data to inform estimates of variables, including unobservable variables like cryptic poaching [16,28,44]. For this study we define unknown fates as those animals that are radio-collared but become lost-to-contact and unmonitored. Most studies involving mortality data make assumptions that unknown fates resemble known fates [16]. When wolves are legally killed, they are always included in known fates and in calculated mortality risk. Treves et al. [16] tested the hypotheses in populations where cryptic poaching occurs, unknown fates will not accurately estimate known fates causing important losses of information producing systematic error. When they corrected estimates of mortality risk of four endangered wolf populations by excluding legal killing from unknown fates, their estimates of poaching risk were higher than government estimates, which assumed known fates would be representative of unknown fates. For example, when estimates for relative risk from other human causes included unknown fates, government reported risk for red wolves was 0.26 – 0.40 lower than corrected estimates [16]. By accounting for all marked animals, *m* (unknown fates) + *n* (known fates), and estimating cryptic poaching, we extract more information for mortality risk than traditional methods of censoring those marked animals that disappeared [16]. We will adapt the above methods to test the hypothesis that censoring or ignoring marked wolves that disappear under-estimates poaching and loses essential information to produce a systematic bias in conclusions about the NENC red wolves.

Here we 1) estimate mortality risk for 5 COD’s (legal, poached, vehicle, nonhuman, and unknown) in the red wolf population from 1987 – 2018, 2) test for association between risk and hunting season, and 3) compare “time to event” for fate unknowns and various CODs. Our analysis adds several years of data not included in prior work and spans a period with a large decline in the wild red wolf population.

## Materials and Methods

### Study area

The red wolf recovery area (RWRA) consists of 6,000 km^2^ of federal, state, and private land in five counties on the Albemarle Peninsula, Northeastern North Carolina (NENC): Beaufort, Dare, Hyde, Tyrrell, and Washington. This area includes four USFWS managed National Wildlife Refuges: Alligator River, Mattamuskeet, Pocosin Lakes, and Swanquarter, a Department of Defense bombing range, and state lands (Fig 1). Land cover types include 40% woody wetlands, 26% cultivated crops, 16% evergreen forest, 5% emergent herbaceous wetlands and other minor (less than 5%) land covers of developed, barren land, deciduous forest, mixed forest, shrub/scrub, herbaceous, and hay/pasture [45]. Elevation ranges between 0-50 m and climate is temperate with four distinct seasons [46].

**Fig 1.**
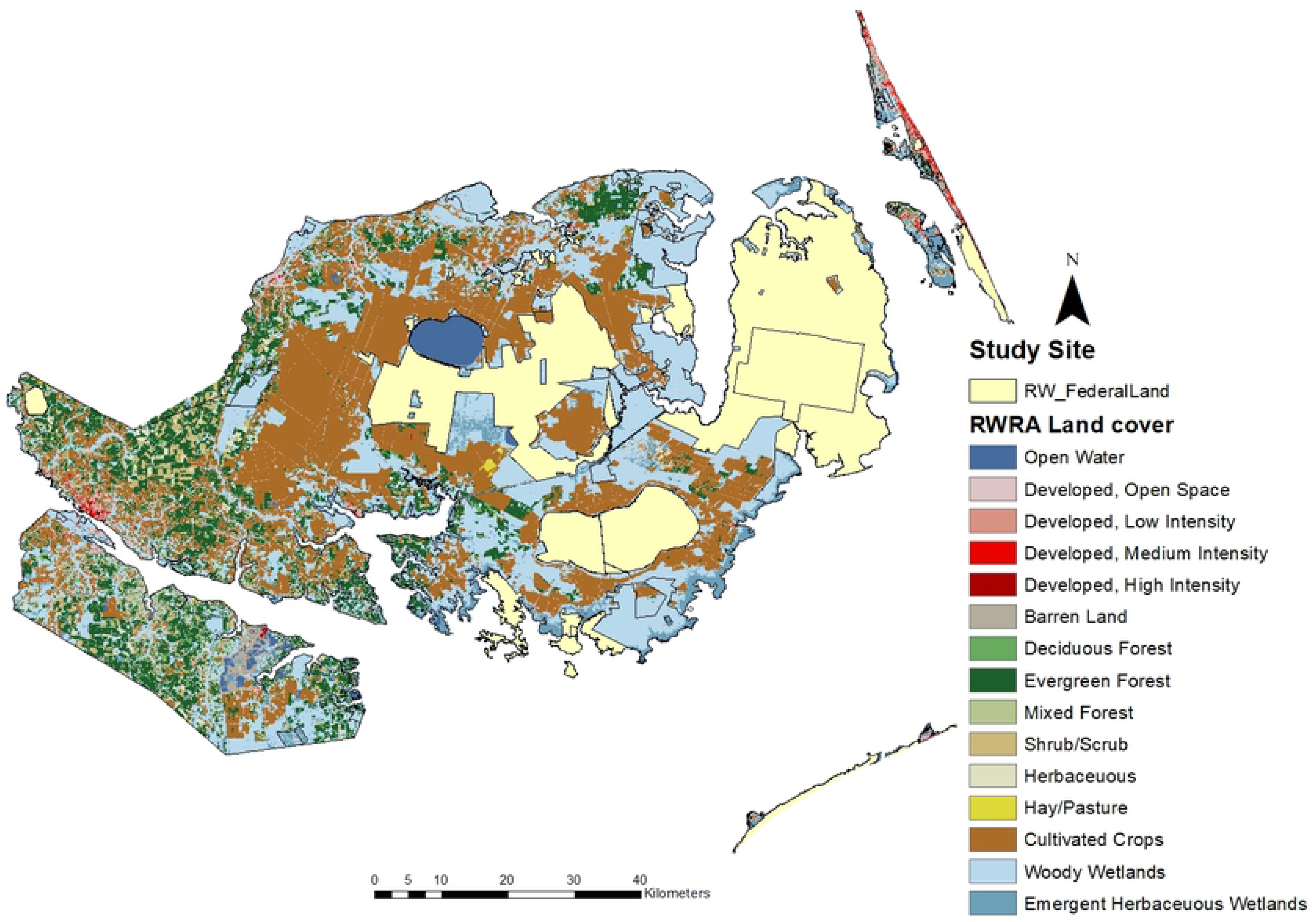
Red wolf recovery area (RWRA) in NENC showing federal and state-owned lands and land cover types. [45].

Red wolves select agricultural habitats over forested areas and transient wolves select edges and roads in these areas more than residents [46]. This use of agricultural lands is in contrast to gray wolves in Wisconsin, which used these types of human landscapes less than expected by chance [17]. Red wolves that maintained stable home ranges between 2009-2011 used 25-190 km^2^ in area, while transients ranged 122-681 km^2^ [46].

### Red wolf sampling

#### USFWS Monitoring

Since reintroduction in 1987, the USFWS estimated abundance of red wolves [22] and maintained a database of all red wolves in the RWRA [47].. Efforts were made to collar every individual, however, radio-collared wolves are not a random sample of all wild red wolves because trapping locations are limited to those areas accessible by USFWS personnel, unknown wolves may have wandered out of the 5-county recovery area (dispersers) or were never caught, and pups (<7.5 months) were not radio-collared for their own safety. Red wolves were captured on federal and state lands as well as private lands with permission from landowners [46]. Following Hinton et al. (2016) we define a red wolf year as October 1 – September 30.

The USFWS database contains information on trapping, tagging, and demographic and spatial information on 810 red wolves including an initial, suspected cause of death (COD), a final, official COD, and necropsy results if performed for each carcass found. These data were collected by the USFWS through field work, reports from trappers, private citizens, and mortality signals from radio-collars [47].

We used population estimates from each year from three sources for red wolf population data: the USFWS database, Hinton et al. (2016), and the 2016 Red Wolf Population Viability Analysis [22,47,48]. Population estimates from these three sources were consistent until 2000 when they began to vary across sources. We do not know why the variation arose, so when the three sources varied, we presented a range of values from the three sources and used the median for analysis.

#### Classifying fates

We analyzed data for 508 radio-collared adult red wolves between the years 1987 – 2018 (63% of all red wolves found in the database; the remaining 37% were pups or uncollared red wolves, which we excluded). The 508 include 393 wolves of known fate and 115 wolves of unknown fate (Table 1). We reclassified USFWS cause of death for known fates into 5 mutually exclusive classes: nonhuman, legal, poached, vehicle, and unknown COD (Table 2). Legal refers to legal removal by USFWS or by a permitted private individual; legal is the only perfectly documented COD in our dataset. All other CODs have different amounts of bias depending on cause, which we refer to as inaccurately documented causes of death, following Treves et al. [17]. Because the ESA made it illegal to kill a listed species except in defense of life [49], we define poaching as any non-permitted killing of a wolf such as shooting, poison, trapping, etc., even if the intended target animal was not a red wolf [17]. We classified “vehicle” separately from poaching because the driver likely did not plan to kill any animal, following [16]. The USFWS listed 12 instances of “suspected foul play”, such as finding a cut collar but no wolf. We classified those as poached, following Hinton et al [22] and Treves et al. [17], because there are only two possibilities with a cut collar; someone removed the collar of a wolf that had already died of another cause, or they killed the wolf, both of which are illegal. Nonhuman causes include mortality related to intraspecific aggression and health related issues such as disease and age. Finally, unknown causes were carcasses whose COD could not be determined. These are important to include in our analyses, as discarding them would overrepresent perfectly reported legal killing [16].

**Table 1.** Annual red wolf population size, COD and FU for collared, adult red wolves.

| Mortality and FU totals for wolf year Oct 1 - Sept 30 |  |  |  |  |  |  |  |
| --- | --- | --- | --- | --- | --- | --- | --- |
|  | Population Est. | Poached | FU | Legal | Collision | Nonhuman | Unknown |
| 1987-88 | 16 | 0 | 0 | 0 | 2 | 2 | 0 |
| 1988-89 | 15 | 0 | 0 | 0 | 1 | 2 | 0 |
| 1989-90 | 31 | 0 | 2 | 0 | 2 | 0 | 0 |
| 1990-91 | 34 | 0 | 0 | 0 | 2 | 4 | 0 |
| 1991-92 | 44 | 0 | 0 | 0 | 0 | 0 | 1 |
| 1992-93 | 67 | 0 | 1 | 0 | 0 | 2 | 0 |
| 1993-94 | 52 | 0 | 1 | 1 | 4 | 4 | 5 |
| 1994-95 | 44 | 7 | 2 | 1 | 1 | 2 | 3 |
| 1995-96 | 52 | 3 | 5 | 1 | 1 | 1 | 3 |
| 1996-97 | 46 | 0 | 3 | 2 | 3 | 2 | 1 |
| 1997-98 | 69 | 1 | 7 | 0 | 2 | 1 | 5 |
| 1998-99 | 90 | 3 | 10 | 2 | 0 | 4 | 2 |
| 1999-00 | 104 | 6 | 4 | 9 | 1 | 0 | 3 |
| 2000-01 | 96-108 | 4 | 3 | 0 | 2 | 1 | 7 |
| 2001-02 | 97-121 | 10 | 7 | 4 | 4 | 4 | 0 |
| 2002-03 | 102-128 | 5 | 1 | 3 | 3 | 7 | 0 |
| 2003-04 | 113-149 | 4 | 5 | 1 | 5 | 5 | 1 |
| 2004-05 | 125-151 | 7 | 7 | 2 | 2 | 4 | 2 |
| 2005-06 | 126-143 | 7 | 5 | 0 | 4 | 3 | 3 |
| 2006-07 | 116-134 | 9 | 7 | 1 | 3 | 1 | 2 |
| 2007-08 | 115-137 | 9 | 7 | 0 | 3 | 3 | 5 |
| 2008-09 | 111-138 | 5 | 8 | 0 | 3 | 1 | 10 |
| 2009-10 | 111-135 | 8 | 5 | 0 | 3 | 4 | 4 |
| 2010-11 | 112-123 | 6 | 4 | 0 | 3 | 5 | 6 |
| 2011-12 | 104-127 | 10 | 6 | 1 | 1 | 0 | 1 |
| 2012-13 | 103-112 | 12 | 4 | 0 | 3 | 1 | 3 |
| 2013-14 | 113-149 | 11 | 3 | 0 | 2 | 3 | 1 |
| 2014-15 | 74 | 11 | 4 | 2 | 0 | 0 | 3 |
| 2015-16 | 45-60 | 3 | 0 | 0 | 2 | 0 | 7 |
| 2016-17 | 20, 25-35 | 6 | 3 | 0 | 0 | 1 | 1 |
| 2017-18 | 19, 23-30 | 2 | 1 | 0 | 3 | 0 | 2 |

<sup>1</sup>Sources: USFWS database, Hinton et al (2016), and the 2016 Red Wolf Population Viability Analysis [22,47,48].

**Table 2.**
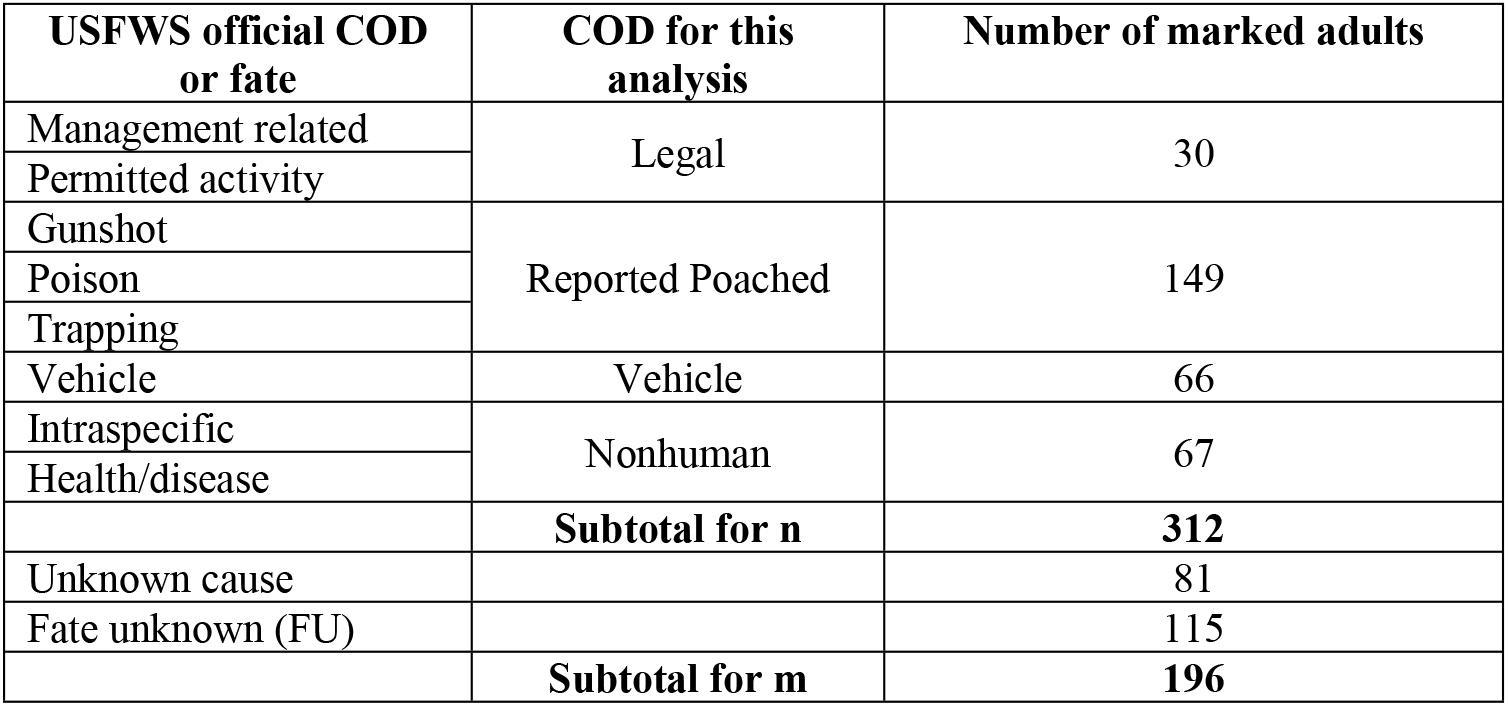
Red wolf fates, our classifications of COD, 1987 – 2018 for 508 radio-collared adults. The variables *n* (known fate subset) estimates the sum of known COD, and *m* (unknown subset) estimates the sum of unknown (FU) and unknown COD. FU are those wolves who were radio-collared but were lost to USFWS monitoring

Fate unknown (FU) wolves were lost to USFWS monitoring and never recovered. Eventually the USFWS stopped monitoring these collars because they could not locate them through aerial or ground telemetry. This could happen if the collar stopped transmitting or if the wolf was killed and the collar was destroyed, referred to as “cryptic poaching” [16,28]. The USFWS assigned their FU date as the date of last contact. We included collared red wolves that might still be alive but unmonitored (FU) at the time of this analysis because they are likely to be dead as of writing and failure to include them would again overrepresent perfectly reported legal killing [16].

If we calculated the risk of each COD as a percent of known fates as is traditional, we would systematically bias against the imperfectly reported causes, disproportionately under-estimate the least well-documented CODs following FU, such as cryptic poaching, and exaggerate the proportion of perfectly reported COD (Legal). Following the method in [16] instead, we estimate risk more accurately by taking into account how many of *m* to reallocate from the unknown COD and FU in Table 2 to our three classes of imperfectly reported CODs (vehicle, poaching, nonhuman). The reallocation step is an estimation procedure with several possible scenarios that apportion different amounts of *m* to cryptic poaching, vehicle, or nonhuman, depending on assumptions that we explain next (Fig 2).

**Fig 2.**
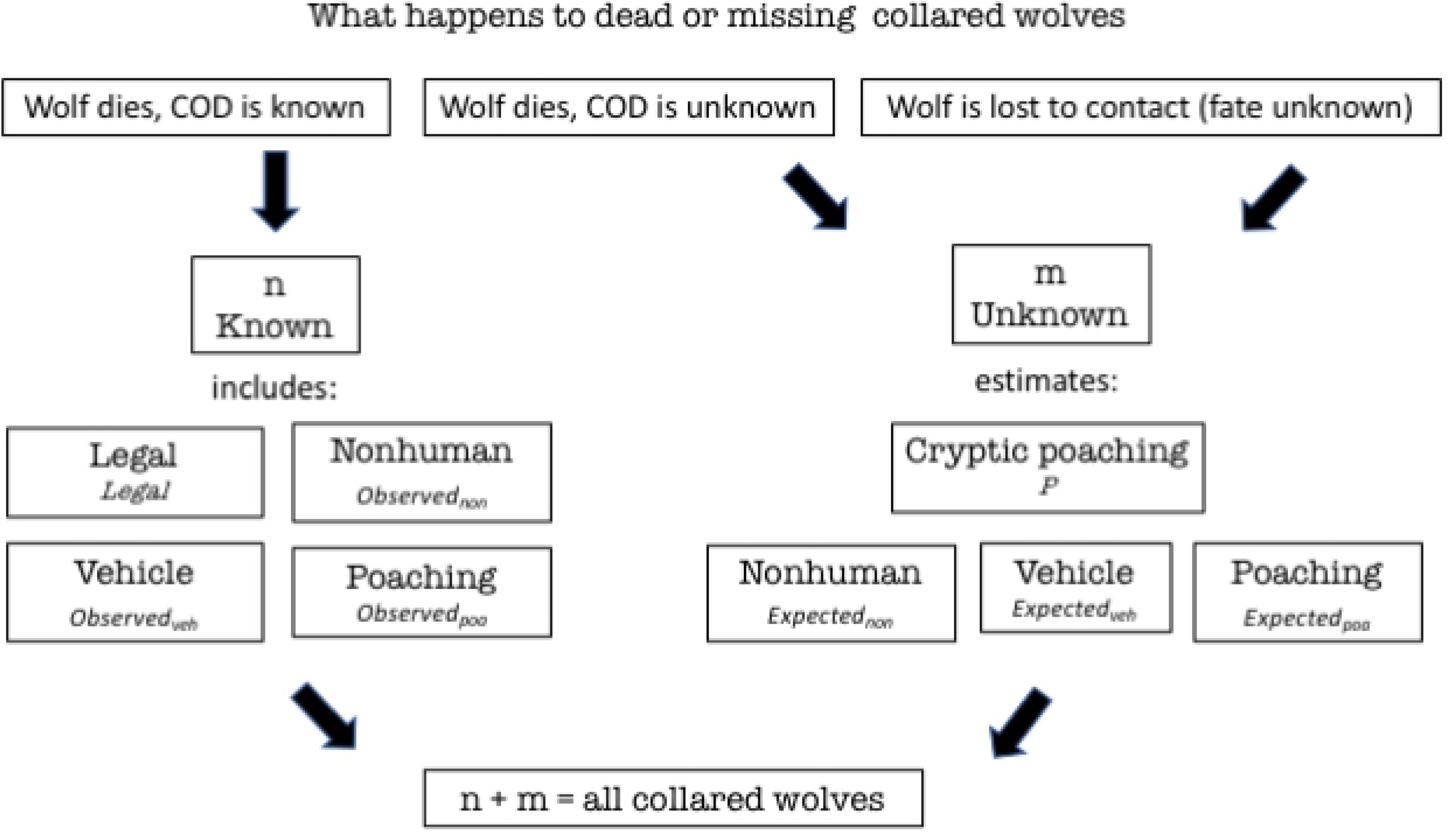
Unknown fates can be estimated. Observed_non_ is the number of marked animals of known fate that died from nonhuman causes. Observed_veh_ is the number of marked animals of known fate that died of vehicle collisions, and Observed_poa_ is the number of marked animals of known fate that died of poaching. Expected_non_ is the number of marked animals of unknown fate expected dead from nonhuman causes, Expected_veh_ is the number of marked animals of unknown fate expected dead from vehicle collision and Expected_poa_ is the number of marked animals of unknown fate expected dead from poaching. P is the estimate of marked wolves dead from cryptic poaching. To account for the possibility of cryptic poaching with transmitter destruction before the wolf meets any other fate, we allocate *m* to the three imperfectly reported CODs. That is why the box for cryptic poaching appears above the others on the right side of the figure.

### Estimating risk from different causes of death

Risk is defined as the proportion of all deaths attributable to a given cause. For example, if five out of ten wolf deaths are the result of poaching, then wolves in that population had a poaching risk of 50%. This is different from mortality rate, which is the rate of individuals dying per unit time, therefore the denominator is all wolves living or dead [17]. To estimate risk accurately, we need the denominator to be all collared red wolves (*n + m*, Fig 2, Table 2). With *n* = 312 wolves with information on COD and *m* = 196 wolves without information on COD (Table 2), the denominator for all risk estimates would be 508. See [16] for a full mathematical description of the method. Below we explain how we adapted it for the red wolf dataset.

First, we calculated the risk of legal killing, in a straightforward manner because they are all known fates perfectly reported by definition and there are none in *m*, the unknown fates portion (Fig 2). Therefore, we had to recalculate the risk posed by the 30 cases of legal killing (Table 1) with the denominator, *n + m*, to estimate the risk of legal killing for all collared red wolves at 5.9% (Table 3). Without this correction, risk of legal killing in Table 1 would be overestimated as 9.6%. When one over-estimates the risk of legal killing, one underestimates all other imperfectly reported CODs because the total must sum to 100%.

**Table 3.** Risk of legal killing among collared red wolves.

|  | Known fates (n) | Unknown fates (m) | Known + Unknown fates (n+m) |
| --- | --- | --- | --- |
| Legal killing | legal/n | 0 | legal/(n+m) |
| Legal killing | 30/312 = 0.096 | 0 | 30/508 = 0.059 |

With the straightforward case of legal killing recalculated as explained above, one can more efficiently explain how to account for the imperfectly reported CODs within *m* (Fig 4).

The unknown fate collared wolves in *m* contain only imperfectly reported CODs but an unknown number of each class of nonhuman, vehicle, and poached. Therefore, we must turn to estimation techniques whose uncertainty produces bounds on our values. The 2 sources for *m* in figure 4 are unknown COD and FU. Poaching that left no trace, such as poisoning in the unknown COD or transmitter destruction in the FU subset, both might contribute to the subcategory of poaching we call cryptic poaching.

Transmitter failure that leads to an animal evading monitoring while the wolf is alive can happen for any of these CODs and account for a portion of *m*. When a transmitter fails and the animal is poached, it is not necessarily cryptic as the transmitter might fail before poaching and the poacher may not have tried to conceal the evidence. However, when the transmitter is destroyed by the poacher (rather than transmitter failure), then we define it as cryptic poaching. Cryptic poaching involves the destruction of evidence [28] distinguishing it from reported and unreported poaching that is present in both *m* and *n*. This changes the way in which we estimate poaching because there is a subset of cryptic poaching in *m* that is never represented in our known fate subset (Fig 2). We assume the destruction of a transmitter occurs earlier than transmitter failure and so when we estimated cryptic poaching, we deducted the estimate of cryptic poaching from *m* first, before we assign the rest of *m* to nonhuman, vehicle, and unreported poaching.

Previous work [16,17] assumed the lower estimate of cryptic poaching was zero. However, we assumed non-zero cryptic poaching in the NENC red wolf population because we had prior information. At least 23 reported poaching incidents in *n* show evidence of attempted and failed cryptic poaching (tampering or damage to the transmitter did not cause its failure, so the USFWS recovered the collar or carcass even though the poacher tried to conceal it). We categorized these 23 as failed cryptic poaching based on the following circumstances; only a damaged collar was found, the dead wolf was found with a damaged collar, or the dead wolf was discovered in a suspicious location (e.g., dumped in a canal, near a beagle that had also been shot also found in that canal, and one was in the same location as another shot wolf). Damaged collars included obvious human tampering such as bullet holes and knife cuts. These 23 deaths include 12 instances categorized by the USFWS as “suspected foul play” and 11 with suspicious circumstances recorded in field notes [47]. Therefore, we inferred that a scenario with zero cryptic poaching was so unlikely as to be discarded. We defined a range of plausible cryptic poaching with the following scenario building.

We used scenario building to triangulate on the probable real value and address uncertainty around a “most likely” outcome as in prior work [50]. Following [16], we used 3 scenarios to estimate cryptic poaching (P). Scenario 1 assumes that poachers who tamper with evidence are equally successful as unsuccessful, so the risk of cryptic poaching in *n* (the 23 wolves described above) is the same as the risk of cryptic poaching in *m*. Therefore, we estimated cryptic poaching as 23*/n* (23/312 = 0.074) meaning P = 0.074 *m =14.4 wolves were successfully concealed for cryptic poaching. We treat this as a lower bound because prior studies of cryptic poaching in wolves have estimated the frequency at 50-69% [17,28] which makes 7.4% seem low.

By contrast our scenario 2 seems like a maximum bound, because we assumed all FU resulted from cryptic poaching but none of the unknown COD did. While this might over-estimate a few cases of transmitter failure, it might under-estimate cryptic poaching in the unknown COD cases that might include killing that was concealed by decomposition or untraceable toxins. Therefore, for scenario 2, P = 115 wolves.

For Scenario 3 we rely on the published estimate of cryptic poaching for Wisconsin wolves, of 46% – 54%. We chose the Wisconsin estimate of two available because the other for Scandinavian wolves would imply P > m, a logical impossibility. The Wisconsin estimate yielded P = 149 wolves.

For all scenarios, we subtracted P from *m* first and then estimated the remainder in *m* as the expected numbers for nonhuman, vehicle, and poaching as follows:

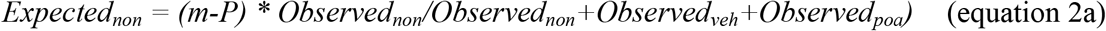

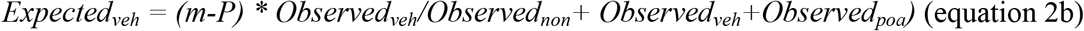

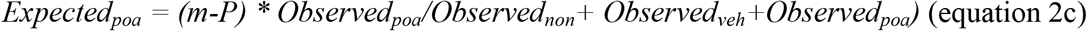

From the three subcategories of poaching, we can estimate total poaching (reported and cryptic).

### Timing of death and disappearance

#### Season

We calculated monthly risk of each COD and used a chi-square analysis to evaluate whether the monthly distributions of CODs differed. We then used a binomial test to evaluate whether CODs during the hunting season (141 days from September 12 – January 31, to include all fall and winter hunting of black bear, deer, and waterfowl) was significantly different from the non-hunting season. Since NC issues nonresident hunting licenses, and several large hunting clubs operate within the five-county RWRA, there were annual influxes of permitted, armed hunters. Using data from 1987-2013, Hinton et al showed a dramatic decrease in survival during the months of October through December [22].

#### Individual survival

We estimated the amount of time individual red wolves spent in the wild from their collaring date to the date of death (with COD) or disappearance (FU) at last contact, all in mean days. We expect a systematic under-estimate of time to FU because monitoring periods were presumably independent of the time that a transmitter stopped, so the last contact always preceded COD. This bias would tend to under-estimate the survival time for the imperfectly reported CODs, especially cryptic poaching relative to all other CODs and other imperfectly reported CODs relative to legal. Addressing this bias and conducting a formal time-to-event analysis was beyond our scope. Therefore, here we visually compared time-to-event across CODs and FU.

We conducted all analysis in STATA IC 15.1 for Mac [51].

## Results

### Estimating risk by cause of death (COD)

From 508 adult radio-collared red wolves, we estimated the risk of legal killing as 0.059 which is less than half of the USFWS estimate of 0.13 [42] and higher than previously published studies of 0.04 – 0.05 [22,24,52]. Relative risk from human COD other than legal ranged from 0.724 – 0.787 depending on the three scenarios for level of cryptic poaching (Table 4), which was 0.026 - 0.044 higher than Treves’ corrected risk estimate using data from 1999 - 2007 [52], and 0.26 – 0.40 higher than government estimates using data from 1999-2007 [42].

**Table 4.** Risk of legal, nonhuman and other human causes of death for known fates (*n*) and unknown fates (*m*) using 3 cryptic poaching scenarios (P from Eqs. 2a – c). Total poaching is the sum of cryptic and reported poaching. Other human is the sum of Vehicle and Total Poaching.

|  |  | Scenario 1, P=14.4 |  | Scenario 2, P=115 |  | Scenario 3, P=149 |  |
| --- | --- | --- | --- | --- | --- | --- | --- |
|  | Risk in *n* | *m* | *n+m* | *m* | *n+m* | *m* | *n+m* |
| Legal killing | 0.096 | 0.000 | 0.059 | 0.000 | 0.059 | 0.000 | 0.059 |
| Nonhuman causes | 0.215 | 0.220 | 0.217 | 0.098 | 0.170 | 0.057 | 0.154 |
| Other human | 0.689 | 0.780 | 0.724 | 0.902 | 0.771 | 0.943 | 0.787 |
| Vehicle | 0.212 | 0.217 | 0.214 | 0.097 | 0.167 | 0.056 | 0.152 |
| Total poaching | 0.478 | 0.563 | 0.511 | 0.805 | 0.604 | 0.887 | 0.635 |
| Cryptic poaching | 0.000 | 0.074 | 0.028 | 0.587 | 0.378 | 0.760 | 0.342 |
| Reported poached | 0.478 | 0.489 | 0.482 | 0.218 | 0.226 | 0.127 | 0.293 |

In scenario 1 with P = 14.4, we estimated total poaching risk at 0.511 with cryptic poaching accounting for 0.055 of total poaching. Scenario 2 with P = 115, our estimate of total poaching was 0.604, which is higher than all previously published estimates (Fig 3) including: 0.33 higher than government estimates and 0.029 higher than Treves 2017. In this scenario, cryptic poaching accounted for 0.626 of total poaching, similar to the Scandinavian estimate [28]. In scenario 3 with P = 149, our estimate of total poaching was 0.635 with cryptic poaching accounting for 0.539 of total poaching as planned by using the Wisconsin estimate.

**Fig 3.**
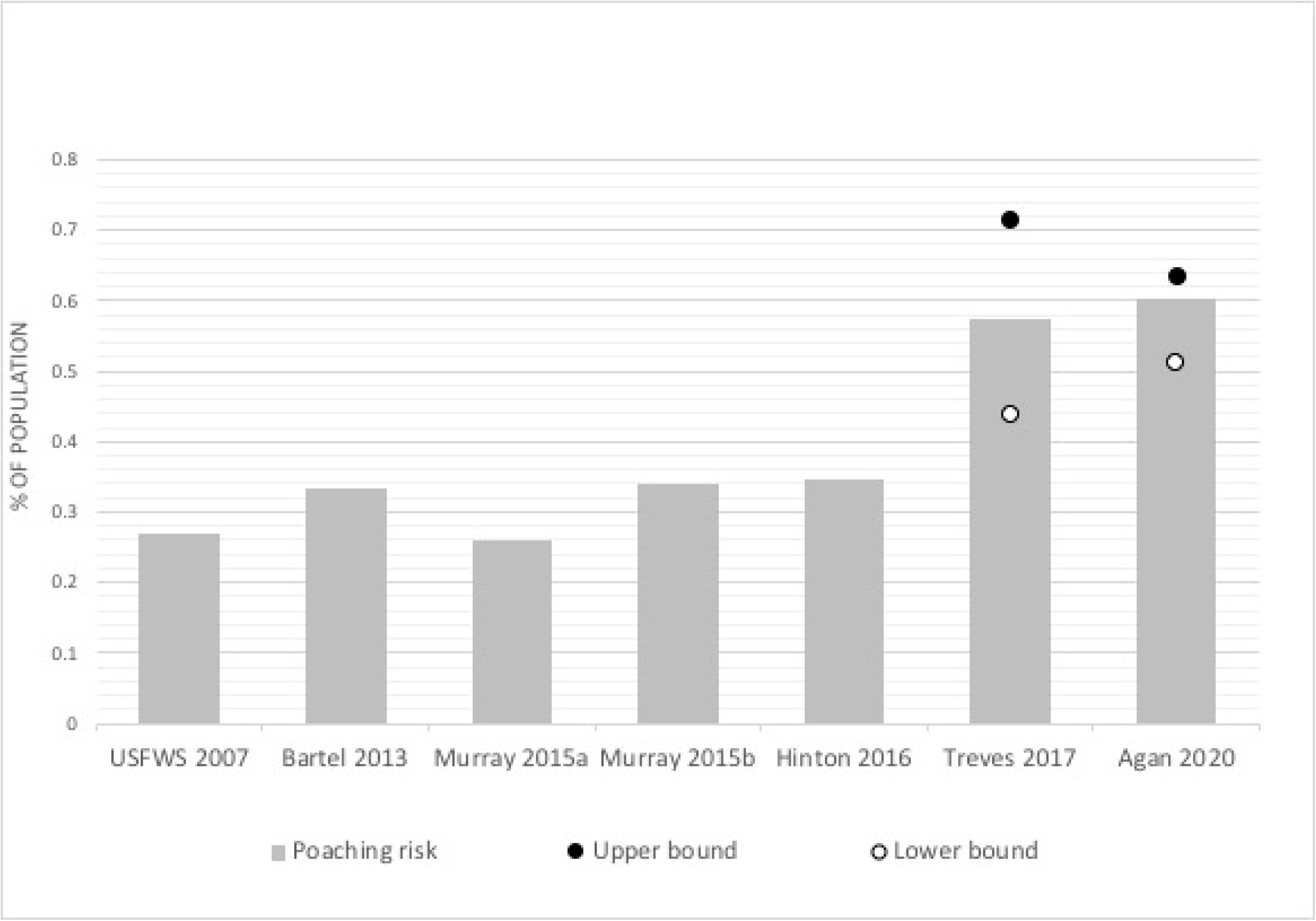
Risk of poaching estimated with different methods from overlapping but different samples of years of the NENC red wolf population. Treves [16] and this study presented 3 scenarios of cryptic poaching with a lower (open circle), middle (bar) and upper (closed circle) bound. Years of data vary by study [22,24,42,52].

### Timing of death and disappearance

#### Season

In our sample of 508 collared, adult red wolves the three months spanning October to December accounted for 61% of all reported poached (October *n*=25, 17%, November *n*=35, 24%, December *n*=29, 20%), which is significantly higher than expected by chance (25%) (Fig 4). The same three months also accounted for 43% of all FU classifications (October *n*=20, 17%, November *n*=12, 10%, December *n*=18, 16%) which is also significantly higher than expected by chance (25%) (χ^2^ = 136.7, df=55, p < 0.001).

**Fig 4.**
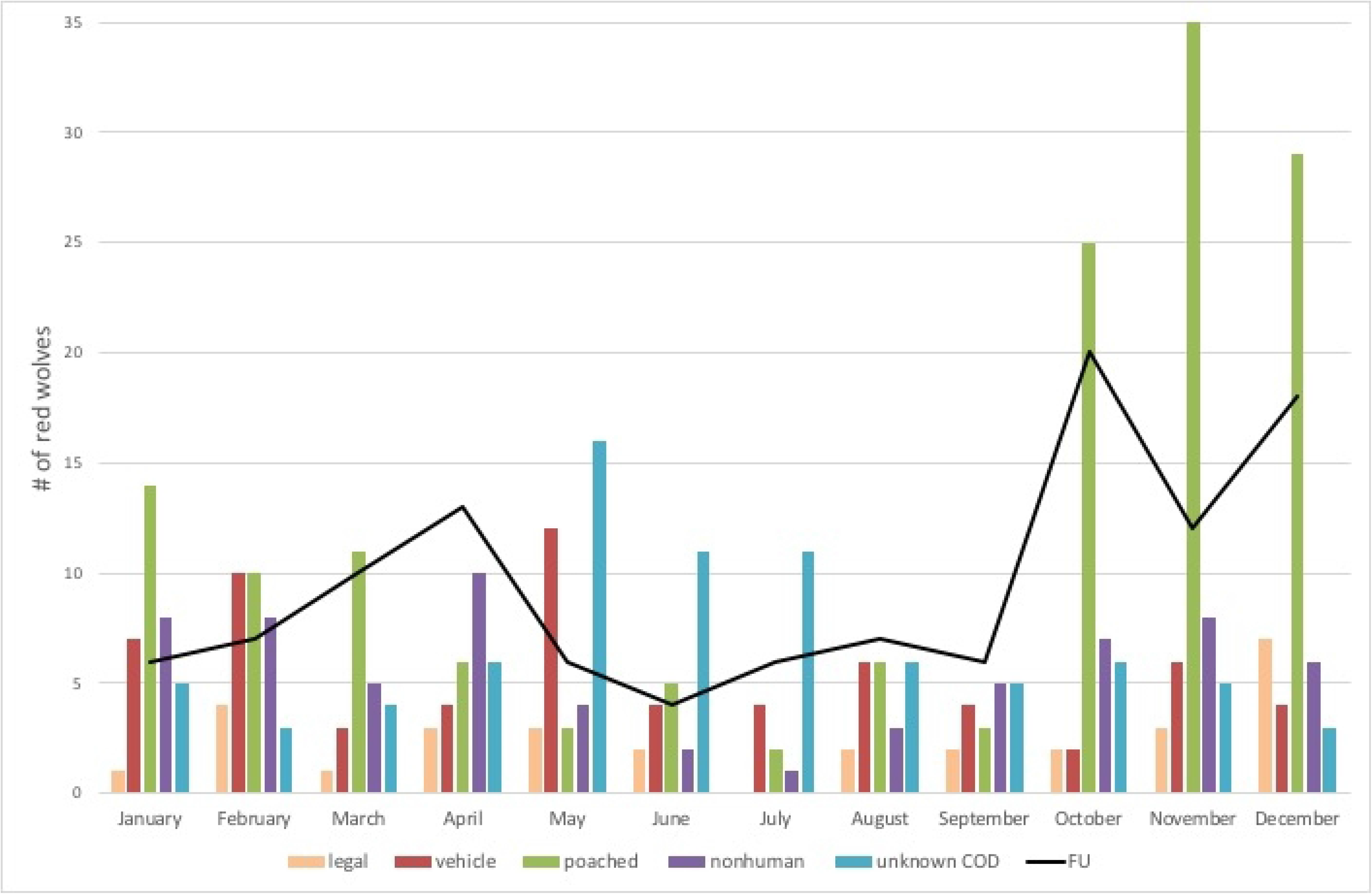
Red wolf mortality each month (bars) compared to fate unknown (FU, black line). FU are represented with a black line because there is more uncertainty with date of disappearance than with date of known deaths. Causes of death include legal, vehicle, reported poached, nonhuman, and unknown cause of death (COD).

We extended our analysis to the full annual hunting seasons of September 12 – January 31 or 141 days. FU and reported poached occurred significantly more often than expected during hunting seasons, (binomial test for FU: expected 38.6%, observed 61 of 115 or 53%, p = 0.002 and reported poached was 104 of 149 or 70%, p < 0.001). Unknown COD was significantly lower than expected during hunting seasons (22 of 81 or 27%, p = 0.04). (Fig 5). There were no significant differences in vehicle, nonhuman, or legal COD.

**Fig 5.**
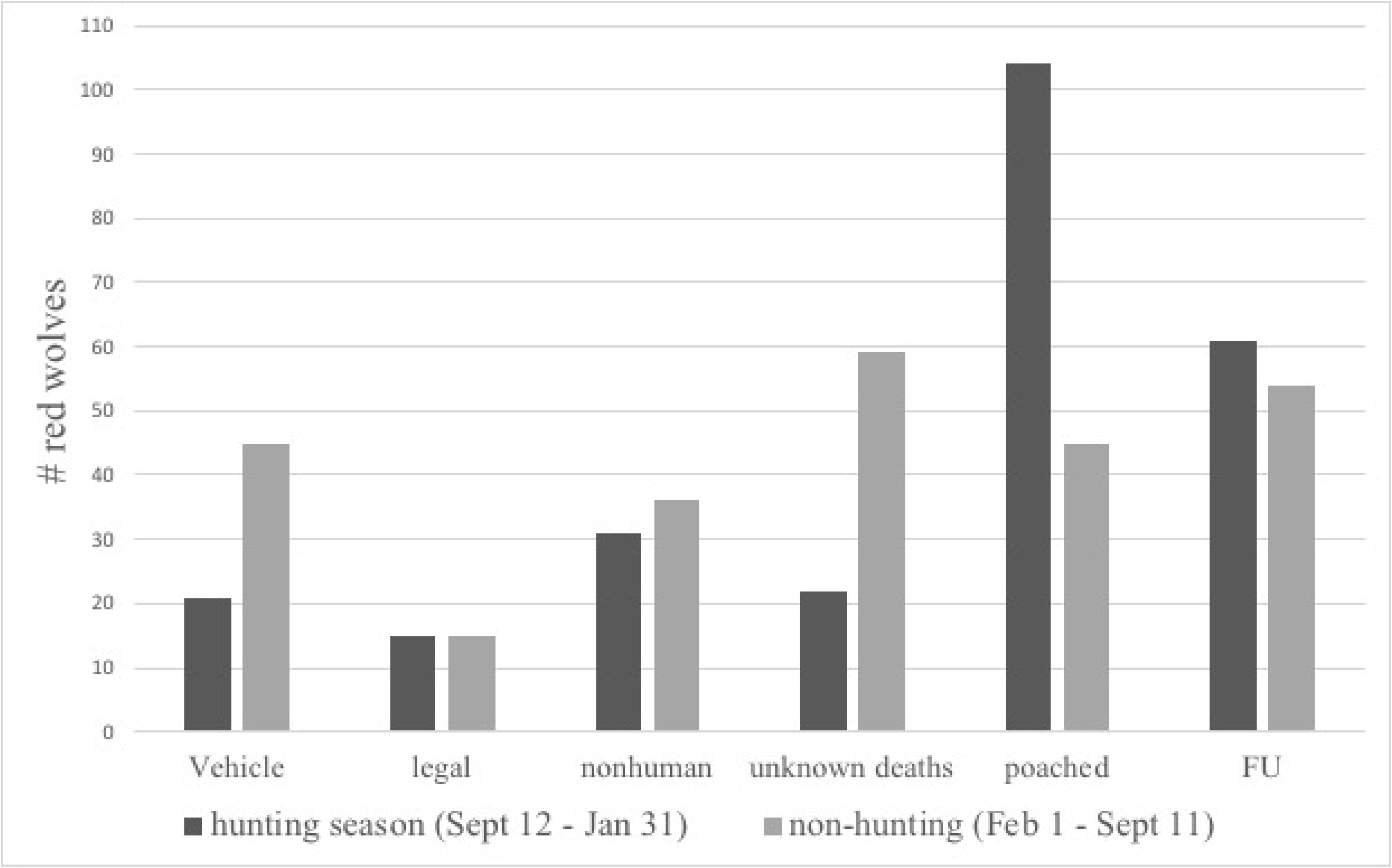
Red wolf deaths and disappearances (FU) by cause during hunting season (dark gray) and non-hunting season (light gray). Hunting season is inclusive of all fall/winter hunting including white-tailed deer, black bear, and waterfowl.

#### Time in the wild

Time after collaring until death or disappearance for 508 adult collared red wolves was 1009 ± 45 days SE (deaths only mean = 1067 ± 52 SE, range 0 – 4,651 days (Fig 6). Nonhuman COD had the longest time in the wild and FU the briefest (Fig. 6), with significant differences between FU and CODs (ANOVA F(5,502) = 5.72, p < 0.001). Out of 115 wolves classified as FU, 92 disappeared 100 - 3471 days after collaring with 16 wolves between 41 – 100 days, 6 wolves between 20 – 40 days, and only one at 9 days after collaring. Time to disappearance and time to death from legal and vehicle were very similar (Fig 6).

**Fig 6.**
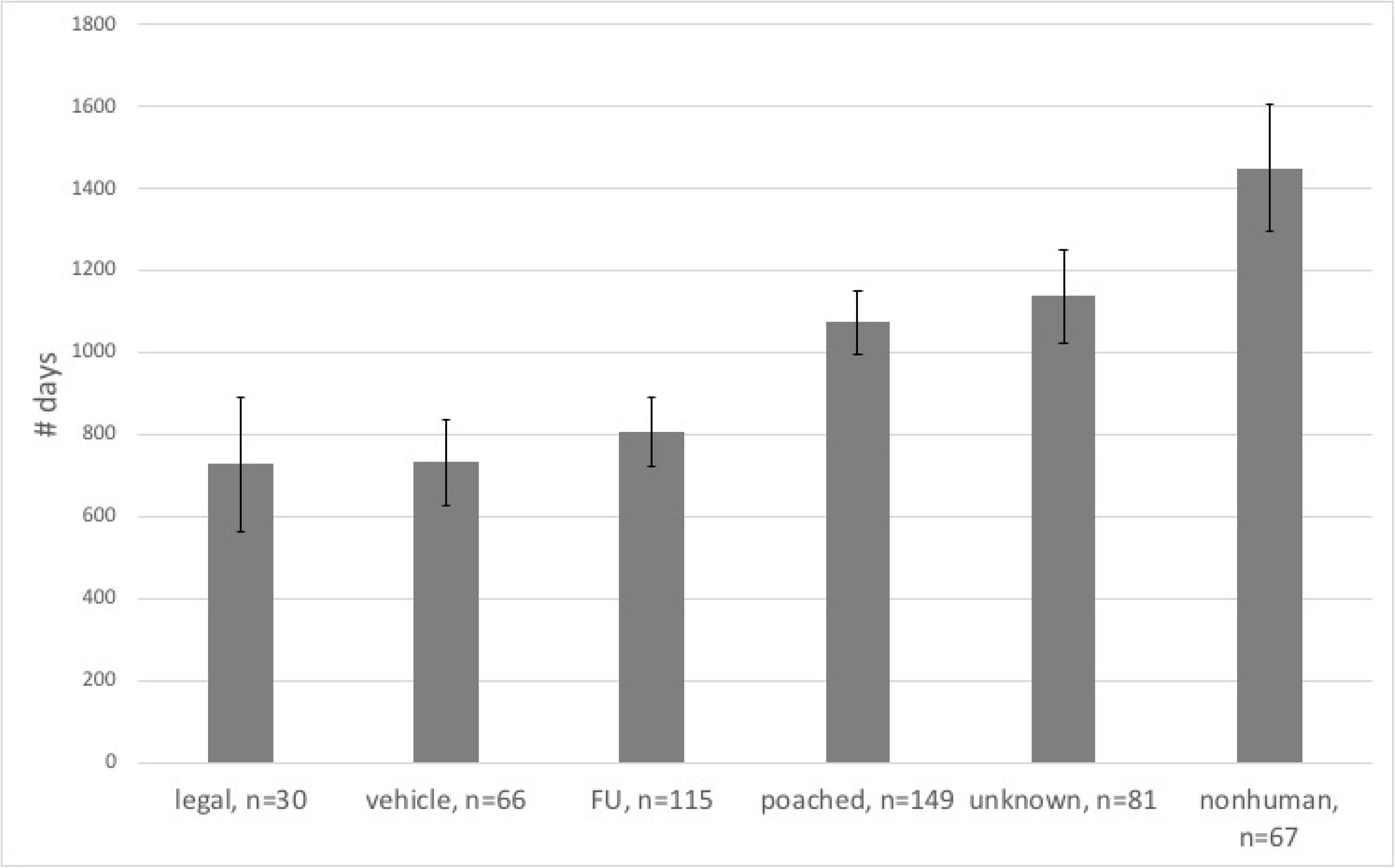
Mean days from collaring to death or disappearance (FU).

## Discussion

Our results for risk of various causes of death (COD), when compared to previous published estimates, support the need for including unknown fates in risk estimates. Legal risk would have been overestimated using only a known fate model, leading to other imperfectly reported CODs being underestimated, first predicted by [16]. By summing known and unknown fates, our risk from anthropogenic causes is higher at 0.78 than previous estimates of 0.45 [52] and 0.61 [36]. The last corrected estimate (conducted with our same method) using data through 2007 [16] was slightly lower than ours at 0.77, suggesting either that uncertainty in both estimates make the two entirely overlapping despite our inclusion of 11 additional years of data or possibly that there has been a slight increase in anthropogenic mortality since that time. Ultimately, anthropogenic mortality reduces the probability of a self-sustaining population, the goal of the USFWS red wolf recovery plan, and could ultimately cause the extinction of the species in the wild in the absence of intervention [53]. Human CODs also violate the Endangered Species Act prohibitions on take.

We estimated total poaching at 0.51 – 0.635 of 508 collard, adult red wolves, with cryptic poaching alone accounting for 0.028 – 0.378 (5-63% of total poaching). Our most conservative estimate for the risk of total poaching at 0.51 is also higher than any previously published risk estimate for red wolves, which ranged from 0.26 [52] to 0.44 [16]. Moreover, our less conservative scenarios estimate cryptic poaching at 0.54 and 0.63, similar to the Wisconsin estimate of 0.50 [16] and the Scandinavian estimate of 0.66 [28]. All three of our red wolf cryptic poaching scenarios lead to very similar total poaching risk, suggesting that regardless of the scenario, wolves in NENC are at higher risk from poaching than any other COD. This supports previous work that suggested poaching might be the cause of decline of this introduced population compared to success in another reintroduced population in Yellowstone [54]. We also suspect poaching of red wolves is higher than any other wolf population measured thus far.

We detected an increase in risk of reported poaching and disappearances during the late fall hunting seasons, similar to other studies that have shown population declines during this same season in red wolves [22], gray wolves [17], and coyotes [55]. Just before fall hunting season, agricultural fields in this region, which make up approximately 30% of the red wolf recovery area [46], are typically cleared of protective cover for wolves [22]. This pattern is quite different from patterns during regulated summer hunting for turkeys from April – May when fields remain covered, vegetation is in leaf, and we found no significant increase in reported poaching or disappearances of red wolves. Although any type of hunting can mean more opportunity for poaching, the lack of an increase in red wolf poaching during the turkey season suggests the hypothesis that other types of hunters are implicated in red wolf poaching. The number of hunting licenses has increased every year in each of the five counties [56] and additional hunters on the landscape could mean more opportunities for poaching. In our study, all other mortality risk for collared, adult red wolves decreased (vehicle, nonhuman, unknown) or remains the same (legal) during legal hunting seasons on other wildlife.

Adult, collared, red wolves also disappeared (FU) more during hunting seasons, suggesting a possible relationship with poaching. This finding is also consistent with the assumption that many FU’s result from cryptic poaching (scenario 2). We estimated FU as cryptic poaching, but many readers will wonder if fate unknown (FU) might not be largely emigrants out of NENC or transmitter failures followed by deaths of other causes in which carcasses were never recovered, rather than cryptic poaching. We draw on research with Wisconsin gray wolves and Mexican wolves [17,57] that provided numerous independent lines of evidence that the majority of FU could not be emigrants nor transmitter failures. First and most importantly, the gray and Mexican wolf studies demonstrated that rates of FU changed with policies on legal killing, which could not plausibly have caused transmitter failures. We cannot see a plausible reason why transmitter failure would explain red wolf FU when it did not for gray and Mexican wolves. Might a difference in ecosystem or latitude play a role in transmitter failure in NENC? Lower temperatures are associated with battery failures, yet the seasonal pattern of FU in NENC matches that of reported poaching, not the annual low temperatures that occur in the months of January through March in NC. Also, average low temperature in NC for this period is between 30 – 40 degrees, while WI averages between 0 – 16 degrees making transmitter failure due to low temperature even less plausible. Also, battery life would seem to play a greater role if FU occurred long after collaring. Contrary to this expectation, FU (808 ± 84 SE days) was much shorter than nonhuman COD (1450 ± 154 SE days) . Second, if FU were largely made up of emigrants, these individuals would likely have died on roads in the densely settled counties beyond the NENC and some of these would have been reported to law enforcement. Even if emigrant collared red wolves escaped the NENC peninsula, some would have been found by citizens with nothing to hide who presumably would have reported their observations to authorities. No such cases are known to us and USFWS data show FU locations clustered amongst other CODs rather than along the edges of the RWRA. Also, dispersal of radio-collared red wolves seems unlikely in an ecosystem with low saturation and abundant, frequent vacancies in territories. Finally, our scenarios encompass non-poaching related explanations for FU and yet total poaching exceeds all other CODs given the known fates and all outcomes we considered.

Some of this poaching, whether cryptic or reported/unreported, is sometimes attributed to mistaken identity. Since red wolves were reintroduced, coyotes have migrated into eastern NC and have been subject to intense shooting and trapping control efforts by regulated hunting [23]. Efforts therefore have been made to limit those occurrences of red wolf killing. Current NC state hunting regulations in the five-county red wolf recovery area restrict coyote hunting to daytime hours only and with a permit [58]. However, this continued allowance of coyote killing reflects a blind spot to the problem of poaching and further efforts to limit this mistaken identity may be necessary. The ESA “Similarity of Appearance” clause allows the Secretary to treat any species as an endangered species if “the effect of the substantial difficulty (to differentiate between the listed and unlisted species) is an additional threat to an endangered or threatened species, and such treatment of an unlisted species will substantially facilitate the enforcement and further the policy of this Act.” (ESA, Sec.4.e). The 1987 amendments to the ESA do not define “knowingly” to include knowledge of an animals species or its protected status, rather it means the act was done voluntarily and intentionally and not because of a mistake or accident [49]. The McKittrick policy which required a perpetrator must have known they were shooting a listed species before they could be prosecuted, was challenged successfully in federal district court [59], then overturned in the appellate court. Newcomer et al. consider Congressional intent in drafting the ESA was clear that harming a listed species would be a crime regardless of the intent or knowledge of the perpetrator, but the McKittrick policy weakens that protection [49].

Our analysis relied on USFWS data for COD and disappearances because we could not verify these independently, which is a limitation of this study. Also, the USFWS has not used GPS collars on red wolves since 2013 and VHF location data includes death locations but not all wolf movements. Previous survival analyses on wildlife populations have not accounted for certain causes of death and disappearance such as policy changes, nor have they been able to model how individual wolves experience policy over time. Traditional survival analyses do not accurately represent marked animals that disappear (FU) since most are censored and not included in outcomes. While survival analysis can model changes in hazard rates, it does not verify why those changes take place such as possible social reasons for increased or decreased poaching. We recommend a time-to-event survival analysis that includes policy period as an intervention since increased poaching appears to be correlated with policy volatility [57].

Although there could be other factors affecting poaching numbers, political volatility [60], and evolving state policy appears to have affected killing of red wolves. For example, North Carolina House Bill 2006, effective January 1, 1995, allowed landowners to use lethal means to take red wolves on their property in Hyde and Washington counties in cases of defense of not only human life (as was always allowed by the ESA regulations) but also threat to livestock, provided the landowner had requested removal by the USFWS. After this policy was enacted, four wolves were shot over the course of Nov 1994 to Dec 1995 [60], the first known to be poached since reintroduction. It seems that both state and federal policy in NC have been less positive for red wolves and may partially explain why the population has declined to near-zero [61]. The USFWS was repeatedly warned about the problem of poaching while there was still time, with a population of 45 – 60 in 2016 [22,48] and beginning in 2017 were notified of the problem of cryptic poaching being underestimated [16]. Those results were shared with the USFWS in public comments in 2018, an official peer review in 2019 and earlier warnings were repeated when comments on red wolves were solicited. It appears there has been negligence or intentional disregard for poaching and the mandates of the ESA.

The USFWS will need to implement favorable policy that prohibits, and interdicts take of red wolves. Killing of red wolves for any reason, other than defense of human life, may actually devalue wolves leading to increases in poaching [29,57]. Rather, programs that reward the presence of wolves can increase their value to local residents. One example of favorable policy related to carnivores was the wolverine program in Sweden, which offered rewards to Sami reindeer herding communities for having reproducing female wolverines on their communal lands [62,63]. Across North America, success of carnivore populations has been linked to legal protections and enforcement [57,64]. Therefore managing wolves successfully, by protecting wolves and encouraging acceptance, might be one way to generate support for their restoration [65]. In NENC policy has changed at both the state and federal level several times since reintroduction began and is currently being reviewed for further changes. This makes it difficult for residents to understand what current regulations are and can lead to confusion about what is allowed or not allowed with regard to killing red wolves. To maintain a positive working relationship with private landowners, USFWS might adapt and enforce policy that is consistent, clear, and protects wolves throughout the entire red wolf recovery area [66] even if it means standing up to illegal actors and communities that condone such law-breaking. Persuading the public to support red wolf recovery might be difficult if landowners resist ESA protections for the animals.

Because approximately 76% of the RWRA is private, and poaching occurs more on private land than public [47], any anti-poaching measure implemented must be proven to work on these lands. Knowledge of wolves’ locations through radio-collaring, trail cameras, and other methods simplify all aspects of management by allowing biologists to locate wolves for any reason including human conflict and should be a part of anti-poaching management. In the past, the USFWS had access to approximately 197,600 acres of private lands through both written and oral agreements. We believe that this has decreased in recent years, both because there are fewer wolves on private land, and because some landowners are no longer as supportive of the program or as willing to allow access, but we cannot quantify the change.

This study suggests aggressive interventions against poaching immediately would be needed if there is going to be any chance the remaining 20 red wolves [61] can create a self-sustaining population. In their most recent 5-year review completed in 2018, the USFWS recommended the red wolf retain its status as endangered under the ESA [67]. While there have been significant changes in the RWRA since reintroduction began, such as the migration of coyotes into the area and rising problems with poaching, the area still retains most of what made it appealing to reintroduction in the first place including low human density and suitable habitat of large areas of woody wetlands, agricultural lands, and protected areas. With the Red Wolf Adaptive Management Plan [68], the USFWS took an aggressive and successful approach to the encroachment of coyotes [68] and should do the same with poaching as mandated by the ESA. Implementing proven practices that prevent poaching or hasten successful reintroduction can reverse the trend of a decreasing NENC red wolf population and once again allow red wolves to thrive, not only in NENC but additional future reintroduction sites.

## Acknowledgments

We thank the USFWS for access to their red wolf database and numerous conversations about the reintroduction program in NENC. We thank Dr. Joseph Hinton from the University of Georgia who provided not only access to his field notes and data but also his guidance and expertise regarding the red wolf program from a research perspective. Thank you to the Carnivore Coexistence Lab at the University of Wisconsin Madison, especially Dr. Francisco J. Santiago-Avila, for their statistical assistance and brainstorming sessions.

